## Supplementary material for "A phylogenetically-restricted essential cell cycle progression factor in the human pathogen *Candida albicans*"

**This PDF file includes:**

Supplementary Text
Figs. S1 to S6
Tables S1 to S5
References

**Supplementary Text**

**Strains, plasmids and primers.** The list of all the yeast strains, primers and plasmids used in this study are mentioned in supplemental tables S3, S4 and S5, respectively. The *E. coli* library harboring CIp10-$P_{TET}$-GTW derivatives was constructed in the TOP10 *ccdB*$^R$ (Invitrogen) strain of *E. coli* (*1*). Other plasmids were propagated in *E. coli* strain DH5α or XL-1 Blue.

*Construction of CSA reporter strain*

The *RFP* was amplified by PCR using primers RFP-PstI-F and RFP-NheI-R from plasmid pNIM1R-RFP (*2*). The PCR fragment was cloned into a TOPO®-TA vector (Thermofisher), digested with PstI and NheI and cloned into the PstI and NheI sites of pTDH3-GFP-URA3 (*3*), yielding pTDH3-RFP-URA3. The *hygB* gene was excised from pAU34-CaHygB (*4*) by BglII+XbaI digest and cloned into the BglII+XbaI double-digested pTDH3-RFP-URA3, to replace the *URA3* marker, yielding pCaTDH3-RFP-HygB. The plasmid pCaTDH3-RFP-HygB was finally modified by a NheI digest and Klenow treatment in order to shorten the extra sequence added at the 3' end because of the cloning steps. The desired $P_{TDH3}$-*RFP-HygB* cassette was PCR amplified from plasmid pCaTDH3-RFP-HygB with oligonucleotides K7_BFP_GFP_Chr4_Right_F and RFP_Insertion_Chr4_Right_Reverse carrying sequences homologous to the genomic DNA located on the right arm of chromosome 4 (Ch4). The PCR product was then transformed in CEC3867 (*5*) yielding CEC5201.

*Generation of a collection* of *C. albicans overexpression strains*

A library of *C. albicans* overexpression strains (1067) was generated using 96-well plates. Briefly, the *E. coli* cultures containing CIp10-$P_{TET}$-GTW derivatives (*6*) were grown in 96-deep well plates containing LB or 2YT medium supplemented with ampicillin (50 or 100 µg/ml). Plasmid minipreparations were carried out in 96-well plates using either Nucleospin™ 96 plasmid core kit (Machery-Nagel™) or the boiling lysis method (*7*). The quality of the isolated plasmids was randomly checked by ethidium-bromide staining following agarose gel electrophoresis. The plasmids were then digested by either StuI (Anza™ 54 Eco147I) or I-sceI

(NEB) depending on whether a *C. albicans* ORF contained a StuI recognition site (*1*). The digested plasmids were precipitated using 3M sodium acetate and 100% ethanol for transformation into the CSA reporter strain (CEC5201), which contains a pNIMX-encoded transactivator to promote the expression from the *TET* promoter (*1*). The *C. albicans* transformation was then carried out in 96 deep-well plates using the lithium acetate method, described previously (*8*). The *C. albicans* transformants were selected for prototrophy and screened by colony PCR using primers PJ88/PJ89 to confirm the integration of the overexpression plasmid at the *RPS1* locus (*1*). The PCR positive transformants were grown in 96 deep-well plates containing YPDU and glycerol stocks of the corresponding strains were prepared.

*Construction of C. albicans overexpression strains*

StuI-digested or I-*Sce*I-digested CIp10-$P_{TET}$-GTW derivatives were used to transform *C. albicans* strains in which the pNIMX transactivator cassette was integrated. The integration of pNIMX at the *ADH1* locus of *C. albicans* was carried out after digesting pNIMX with KpnI and ApaI, and confirming the transformants by PCR using primers PJ86/PJ87 (*1*). The *C. albicans* transformants harboring the overexpression plasmid at the *RPS1* locus were screened by PCR using primers PJ88 and 89.

*C-terminal tagging of Tub4 with fluorescent proteins*

GFP-tagged Tub4 expressing strains were constructed by using the plasmid pTub4-GFP-His. Briefly, the 3' coding region of Tub4 without the stop codon was amplified from the *C. albicans* (SN148) genome using primers LS39FP/LS39RP and cloned into the SacII and SpeI sites of pBSGFP-His (*9*). The resulting plasmid pTub4-GFP-His was confirmed using restriction analyses, digested with PacI and was used to transform the *C. albicans* strains. The correct *C. albicans* transformants were screened by fluorescence microscopy.

Epitope tagging of Tub4 with mCherry was carried out using the plasmids pTub4-mCherry-Arg4 or pTub4-mCherry-Nat. To construct pTub4-mCherry-Arg4, the C-terminus of Tub4 was released from pTub4-GFP-His following digestion with SacII and SpeI and subsequently cloned into the SacII and SpeI sites of pRFP-Arg4 (*10*). The *E. coli* clones were confirmed by restriction analyses. The plasmid pTub4-mCherry-Arg4 was partially digested with PacI for transforming *C. albicans* strains. The transformants obtained were selected for prototrophy and screened by fluorescence microscopy. The plasmid pTub4-mCherry-Nat was constructed as follows: the mCherry coding gene was amplified from CaADH1pyEmRFP (*11*) using the primers SR149/SR150 and cloned into the SpeI and SmaI sites of pBSNAT (*12*). The mCherry-NAT containing plasmid was then digested by SpeI and KpnI and the mCherry-NAT fragment was cloned into the SpeI and KpnI sites of pTub4-GFP-His. The resulting plasmid pTub4-mCherry-Nat was verified by restriction analyses and used to transform *C. albicans* strains after PacI digestion. The correct *C. albicans* transformants were screened by fluorescence microscopy.

*C-terminal tagging of Tub1 with mCherry*

The 3' coding region of Tub1 without the stop codon was amplified from *C. albicans* (SN148) genome using the primer pairs PJ77/PJ88 and cloned into the SacII and SpeI sites of pRFP-Arg4 (*10*). The resulting plasmid pTub1-mCherry-Arg4 was confirmed using restriction analyses and digested with XbaI for transforming the *C. albicans* strains. The correct *C. albicans* transformants were screened by fluorescence microscopy.

*C-terminal tagging of Cse4 with TAP*

The *CSE4* ORF (without the stop codon) along with the *TAP* tag was PCR amplified from the *C. albicans* strain CAKS102 (*13*) using the primers NV241/NV242 and cloned into the SalI and ApaI sites of pMad2-2 (*14*). The desired plasmid pCse4-TAP-Leu was verified by restriction analyses and linearized by XhoI for transforming the *C. albicans* strains. The transformants were selected for prototrophy and confirmed by western blot analysis.

*Construction of bub2 null mutant*

Both the alleles of *BUB2* were deleted using the *SAT1* flipper cassette (pSFS2a) (*15*). To delete the first allele, upstream (US) and downstream (DS) sequences of *BUB2* were amplified from the *C. albicans* (SN148) genome using primers PJ110/PJ111 and PJ112/PJ113, respectively. The US and DS sequences were then cloned in pSFS2a as KpnI/XhoI and SacII/SacI fragments, respectively, to obtain the plasmid pBub2del#1. The *E. coli* clones were verified using restriction analysis. The desired deletion cassette was transformed into the *C. albicans* strains after digesting pBub2del#1 with KpnI and SacI. The *C. albicans* transformants were screened for correct chromosomal integration by PCR using the primer pair PJ3/PJ116. The correct transformants were grown in YPM (1% yeast extract, 2% peptone, 2% maltose) medium supplemented with uridine (0.1µg/ml) overnight and plated on YPMU agar to recycle the *SAT1* marker (*15*). The single colonies obtained on YPMU agar were replica plated on YPDU and YPDU with nourseothricin (100 µg/ml). Nourseothricin-sensitive colonies, obtained because of *SAT1* recycling, were reconfirmed for *SAT1* eviction and *BUB2* first copy deletion by PCR using primers PJ110/PJ113 and selected for subsequent transformation experiments to delete the remaining *BUB2* allele.

To delete the second allele of *BUB2*, the DS sequence of *BUB2* in pBub2del#1 was replaced with the 3' coding region of *BUB2*. For this, the 3' coding region of *BUB2* was amplified from the *C. albicans* (SN148) genome using primers PJ114/PJ115 and cloned into the SacII and SacI sites of pBub2del#1. The resulting plasmid pBub2del#2 was verified using restriction analyses and was used to transform the *BUB2* heterozygous null strain after digesting pBub2del#2 with KpnI and SacI. The transformants obtained were grown in presence of nourseothricin (100 µg/ml) and screened for the integration of pBub2del#2 deletion cassette by PCR using the primers PJ117/PJ118. The desired PCR positive transformants were reverified for *BUB2* first copy deletion using the primers PJ110/PJ113. The resulting nourseothricin-resistant *bub2* mutants were grown in YPMU overnight and plated on YPMU agar to recycle the *SAT1* marker (*15*). The single colonies obtained on YPMU agar were replica plated on YPDU and YPDU with nourseothricin (100 µg/ml). Nourseothricin-sensitive colonies were selected for subsequent experiments.

*Construction of csa6 conditional mutant*

The first allele of *CSA6 (ORF19.1447)* was deleted using the *SAT1* flipper cassette (pSFS2a) (*15*) and the second allele was placed under the control of regulatable *MET3* promoter (*16*). To delete the first allele, the US and DS sequences of *CSA6* were amplified from the *C. albicans* (SN148) genome using primers PJ95/PJ96 and PJ97/PJ98, respectively. The US and DS sequences were then cloned in pSFS2a as KpnI/XhoI and SacII/SacI fragments, respectively, to obtain the plasmid pCsa6del. The *E. coli* clones were confirmed using restriction analysis. The desired deletion cassette was transformed into *C. albicans* strains after digesting pCsa6del with KpnI and SacI. The *C. albicans* transformants were screened for correct chromosomal integration by PCR using the primer pair PJ3/PJ99. The correct transformants were grown in YPMU overnight and plated on YPMU agar to recycle the *SAT1* marker (*15*). Nourseothricin-sensitive colonies, obtained because of *SAT1* recycling, were selected for subsequent transformations to inactivate the remaining wild-type allele of *CSA6*.

To replace the promoter of the second allele with the *MET3* promoter (*16*), the 5' coding region of *CSA6* including the start codon was amplified from the *C. albicans* SN148 genome using primers PJ93/PJ94 and cloned into the BamHI and PstI sites of pCaDis (*16*), generating the plasmid pCsa6-Met3-Ura. The *E. coli* clones were confirmed using restriction analyses. The plasmid pCsa6-Met3-Ura was linearized using BstBI and was used to transform *C. albicans* strains in which the first copy of *CSA6* was deleted. The resulting conditional mutants were screened for correct genomic integration by PCR using the primer pair PJ91/PJ95.

To generate the plasmid pCsa6-Met3-His, a *HIS1* fragment from pGFP-HIS (*9*) was obtained after digesting pGFP-HIS with EcoRI and was cloned into the EcoRI site of p1447-Met3-His. The *E. coli* transformants were screened and desired clones were validated by restriction analysis.

*Epitope tagging of Csa6*

The C-terminus of *CSA6* was tagged with either TAP or mCherry. To express TAP-tagged Csa6 from the native promoter or the *MET3* promoter, the 3' coding region of *CSA6* without the stop codon was amplified from the *C. albicans* SN148 genome using the primer pair PJ108 (containing the BglII restriction site) and PJ109 and cloned into the BamHI and PacI sites of pFA-TAP-*ARG4* (*17*). The *E. coli* clones were confirmed by both restriction analyses and Sanger sequencing. The resulting plasmid p1447-TAP-Arg was linearized using BamHI for single-site integration into the *C. albicans* genome. The *C. albicans* transformants were screened for correct genomic integration by PCR using the primer set NV34/TEJ13 and western blot analysis.

To express Csa6TAP from the $P_{TET}$ promoter, a fragment containing the coding region of *CSA6* along with the *TAP* tag was amplified from CaPJ180 using primers PJ127/PJ128 and cloned into the EcoRV site of pCIp10-$P_{TET}$-GTW (*1*). The resulting plasmid CIp10-$P_{TET}$-1447TAP was confirmed using restriction analyses and Sanger sequencing and was digested by StuI for transforming the *C. albicans* strains. The correct *C. albicans* transformants were screened by PCR using primers PJ88/PJ89 and western blot analyses.

To tag the C-terminus of Csa6 with mCherry, the 3' coding region of *CSA6* without the stop codon was amplified from the *C. albicans* SN148 genome using the primer pair TEJ1/TEJ2 and cloned into the SacII and SpeI sites of pRFP-Arg4 (*10*). The *E. coli* clones were confirmed by both restriction analyses and Sanger sequencing. The resulting plasmid pCsa6-mCherry-Arg was linearized using BstB1 for single-site integration into the *C. albicans* genome. The correct *C. albicans* transformants were confirmed by PCR using primers TEJ13/TEJ14 and analyzed by fluorescence microscopy. The functionality of the mCherry-tagged Csa6 was determined by tagging the only copy of *CSA6* in a heterozygous null mutant (CaPJ209, *csa6/CSA6*) with mCherry. The resulting strain CaPJ117 (*csa6/CSA6-mCherry*) was viable and did not show any growth defect.

*Construction of SOL1 overexpression mutant*

To overexpress Sol1, an extra copy of *SOL1* under the $P_{TET}$ promoter was integrated at the *RPS1* locus (*1*). For this, the complete ORF sequence of *SOL1* including the start and the stop codon was PCR amplified from the *C. albicans* (SN148) genome using primers PJ119/PJ120 and cloned into the EcoRV site of CIp10-$P_{TET}$-GTW (*1*) resulting in plasmid CIp10-$P_{TET}$-SOL1. The correct *E. coli* clones were screened using restriction analyses and verified by Sanger sequencing. The plasmid CIp10-$P_{TET}$-SOL1 can be linearized by StuI for *C. albicans* transformation. The *C. albicans* transformants were confirmed by PCR using primers PJ88/PJ89.

*Epitope tagging of Sol1*

The expression level of Sol1 from the native promoter and $P_{TET}$ promoter was compared by tagging the C-terminus of *SOL1* with TAP. For TAP tagging Sol1 under its own promoter, the 3' coding region of *SOL1* without the stop codon was amplified from the *C. albicans* (SN148) genome using the primer pair PJ141/PJ142 and cloned into the BamHI and PacI sites of pFA-TAP-*His1* (*16*). The *E. coli* clones were confirmed by both restriction analyses and Sanger sequencing. The resulting plasmid pSol1-TAP-His was linearized using XbaI for single-site integration into the *C. albicans* genome. The *C. albicans* transformants were screened for correct genomic integration by PCR using the primer set PJ119/PJ128 and western blot analysis.

To express Sol1TAP from $P_{TET}$, a fragment containing the coding region of *SOL1* along with the *TAP* tag was amplified from CaPJ216 using primers PJ119/PJ128 and cloned into the EcoRV site of CIp10-$P_{TET}$-GTW (*1*). The resulting plasmid CIp10-PTET-SOL1TAP was confirmed using restriction analyses and Sanger sequencing. The plasmid CIp10-PTET-SOL1TAP was linearized by StuI for transforming the *C. albicans* strains and the transformants were screened by PCR using primers PJ88/PJ89 and western blot analyses.

*Construction of GFP-tagged strains of Tem1* and *Spc110*

Tem1 and Spc110 were tagged C-terminally with GFP. For this, 3' coding region of Tem1 or Spc110 without the stop codon was amplified from the *C. albicans* (SN148) genome using the primer pairs PJ121/122 and PJ106/PJ107, respectively and cloned into the SacII and SpeI sites of pBSGFP-His (*9*). The resulting plasmids (i) pTEM1-GFP-His was confirmed by both restriction analyses and Sanger sequencing (ii) pSpc110-GFP-His was validated by restriction analyses.

The plasmid pTEM1-GFP-His was propagated in the *dam⁻/dcm⁻* strain of *E. coli* (C2925) obtained from NEB and digested with BclI for transforming the *C. albicans* strains. The correct *C. albicans* transformants were screened by PCR using primers PJ123/PJ124 and fluorescence microscopy.

The plasmid pSpc110-GFP-His was linearized with NheI or NsiI for single-site integration into the *C. albicans* genome. The *C. albicans* transformants were screened by fluorescence microscopy.

*Expression of C. dubliniensis Csa6 in C. albicans*

The C-terminus of *C. dubliniensis* Csa6 *(Cd36_16290)* was tagged with GFP and ectopically expressed in *C. albicans* using the plasmid pCdCsa6-GFP-ARS2. For this, the complete ORF of *CdCSA6* (without the stop codon), along with its promoter region was PCR amplified from the genome of Cd36, a *C. dubliniensis* clinical isolate (*12*), using the primers VS5/VS6. The *GFP* tag was amplified from pTub4-GFP-His using the primers VS7/VS8. An overlap PCR of the two fragments was then set up using the primers VS5/VS8. The resulting ~3.6 kb long fragment containing the GFP-tagged *C. dubliniensis CSA6* under its own promoter was cloned into the XbaI and PstI sites of pARS2 (*9*). The plasmid, pCdCsa6-GFP-ARS2, obtained was verified by restriction analyses and transformed into the *C. albicans* strains. The transformants were selected for prototrophy and screened by fluorescence microscopy. As ARS plasmids are highly unstable in *C. albicans*, we used the large transformant colonies, obtained as a result of an integrative transformation of pARS2 (*18*) and retained the auxotrophic marker (*URA3)* even in the absence of any selection pressure, for all our assays.

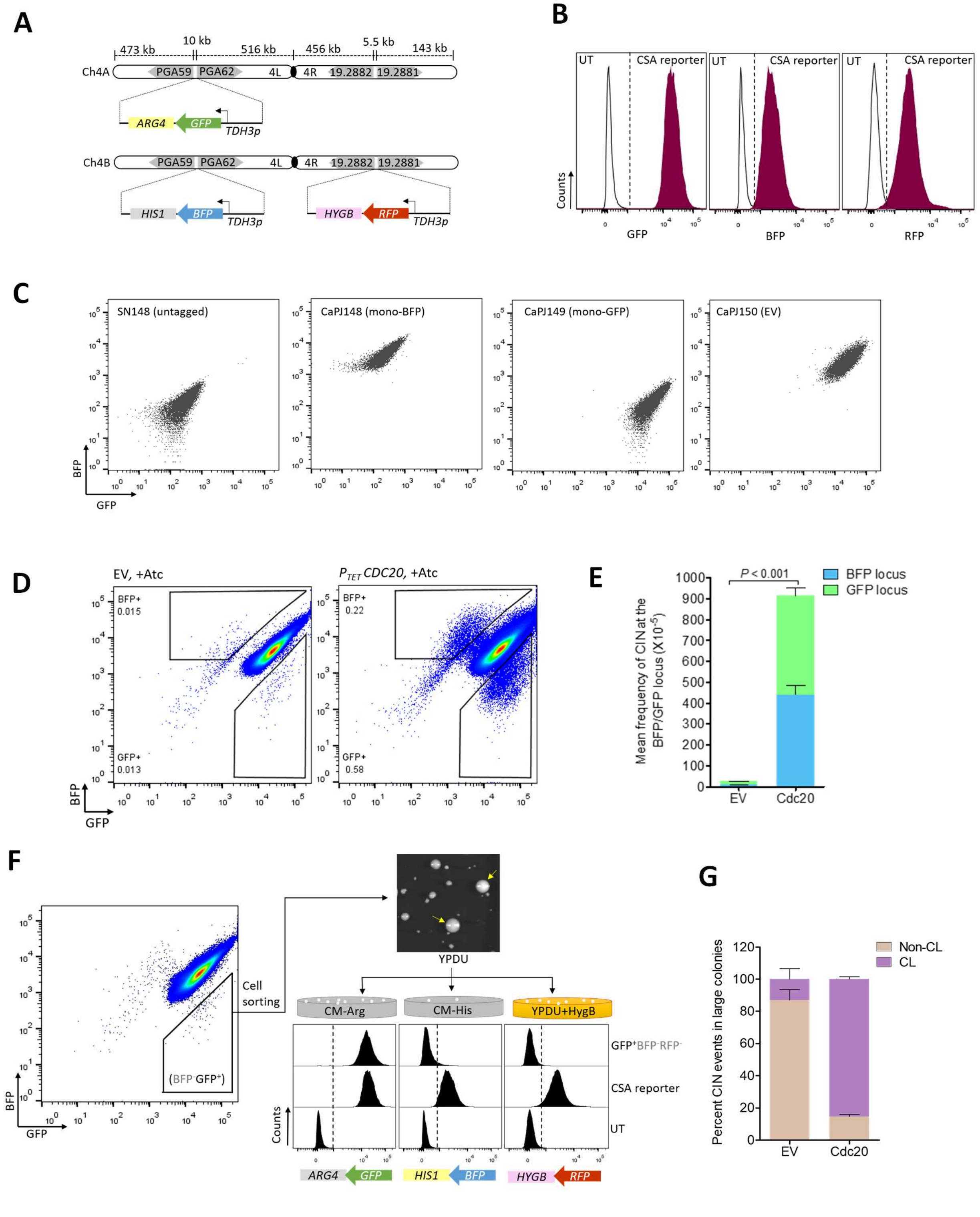


**Fig. S1. The CSA reporter system for detecting chromosome instability (CIN) in *C. albicans*. (A)** A line diagram of Chromosome 4 (Ch4) in the CSA reporter (CEC5201). As indicated, BFP/GFP-expressing cassettes are present on the left arm of both the homologs of Ch4 in the *PGA59-PGA62* intergenic region, while the RFP-expressing cassette is present on the right arm of Ch4 in the *ORF19.2882-ORF 19.2881* intergenic region. Expression of BFP, GFP or RFP is under control of the *TDH3* promoter and is associated with respective selectable markers, as mentioned in the diagram. **(B)** Histograms showing fluorescence intensity measurements of an untagged strain (SN148) and the CSA reporter strain (CEC5201) by flow cytometry. The CSA reporter strain on the *right* exhibits higher fluorescence intensity for GFP, BFP and RFP laser than the untagged (UT) strain. **(C)** Flow cytometric analysis of the BFP/GFP marker in various control strains as indicated. The strains were grown in YPDU medium overnight and analyzed by flow cytometry. Approximately 10,000 events are displayed. **(D)** Detection of chromosome instability in the *CDC20$^{OE}$* strain (CaPJ151). *Left*, a representative BFP/GFP density plot of EV in presence of anhydrotetracycline (Atc) (3 µg/ml), an inducer of $P_{TET}$. The proportion of $BFP^{+}GFP^{-}$ and $BFP^{-}GFP^{+}$ cells in the EV indicates the intrinsic instability of Ch4 in *C. albicans*. *Right*, a representative BFP/GFP density plot of the *CDC20$^{OE}$* strain in presence of Atc (3 µg/ml). **(E)** Quantitation of the mean frequency ($x10^{-5}$) of CIN at the BFP/GFP locus in CaPJ150 (EV) and CaPJ151 (*CDC20$^{OE}$*); *N*=3. Unpaired *t*-test, one-tailed, *P*-value shows a significant difference. **(F)** Schematic illustrating the workflow to differentiate CL from non-CL events. A representative flow cytometry density plot is shown as a reference. $BFP^{-}GFP^{+}$ cells were sorted and plated on YPDU agar. Large colonies (arrow marked in yellow) were tested for the presence of selectable markers, *ARG4*, *HIS1* and *HYGB* by replica plating, followed by flow cytometry analysis to monitor the presence of the associated fluorescent proteins (GFP, BFP and RFP). The fluorescence intensity profile of an *ARG4* resistant colony which had lost *HIS1* and *HYGB* ($GFP^{+}$ $BFP^{-}RFP^{-}$) is shown as an example. The concomitant loss of *BFP-HIS1* and *RFP-HYGB* indicates that the entire Ch4B is lost. **(G)** Analysis of the marker genes, *ARG4*, *HIS1 and HYGB* by replica plating $BFP^{-}GFP^{+}$ colonies of EV (CaPJ150) and *CDC20$^{OE}$* strain (CaPJ151); *N*=3 with ≥100 colonies for each *N*.

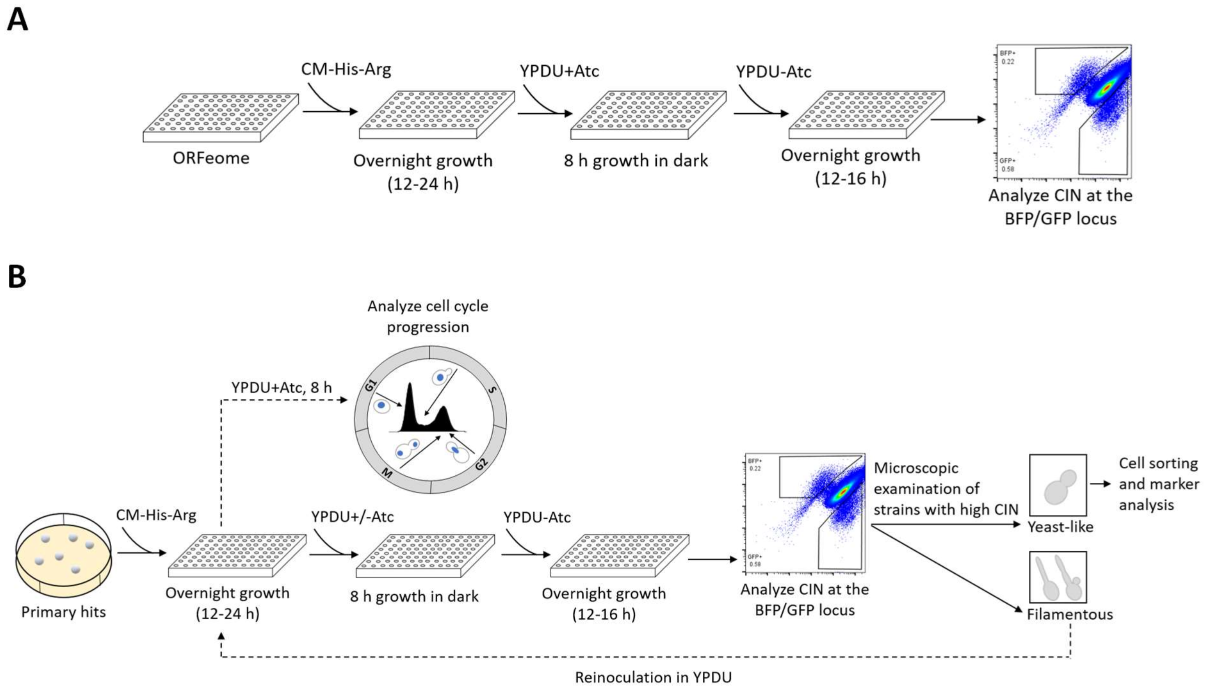


**Fig. S2. Primary and secondary screening of the *C. albicans* overexpression library. (A)** Flow chart illustrating the steps of the primary screen. Briefly, overnight grown cells were induced for 8 h in presence of Atc (3 µg/ml), allowed to recover overnight in a rich medium without Atc, diluted in 1x PBS, and analyzed for BFP/GFP marker by flow cytometry (~$10^6$ cells). Gates were defined for the $BFP^+GFP^-$ and $BFP^-GFP^+$ populations in the EV (CaPJ150) and applied to all other over-expression mutants (1067). Mutants were selected if the frequency of CIN at the BFP/GFP locus was two-fold higher than the frequency in the EV. **(B)** Flow diagram illustrating the steps of the secondary screen. The overexpression mutants identified from the primary screen (23 out of 1067) were induced for 8 h in presence or absence of Atc (3 µg/ml), allowed to recover overnight in a rich medium without Atc and analyzed for the loss of BFP and GFP by flow cytometry. Mutants were selected if they exhibited two-fold higher rate of CIN at the BFP/GFP locus in three biological replicates as compared to the EV and further analyzed for any morphological transition by microscopy. Overexpression mutants with yeast-like morphology were analyzed by cell sorting and marker analysis to determine the molecular mechanism (CL or non-CL) leading to CIN. Overexpression mutants exhibiting polarized growth were regrown, induced for 8 h in presence of Atc (3 µg/ml) and analyzed for cell cycle progression by microscopy or flow cytometry.

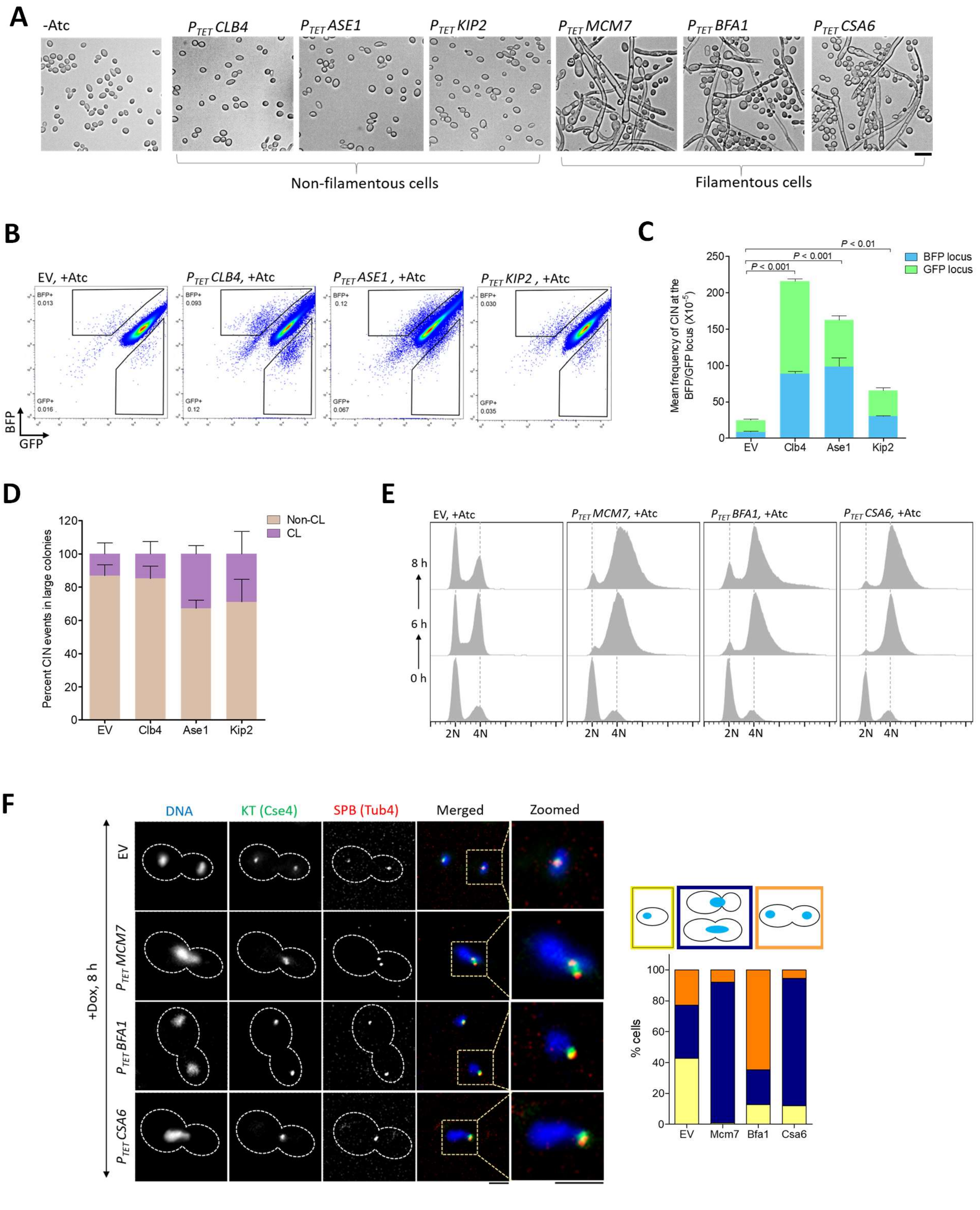


**Fig. S3. CIN associated with overexpression of *CSA* genes is regulated via distinct mechanisms. (A)** Bright-field micrographs of the six overexpression strains, *CSA1*$^{CLB4}$

(CaPJ152), *CSA2*$^{ASE1}$ (CaPJ153), *CSA3*$^{KIP2}$ (CaPJ154), *CSA4*$^{MCM7}$ (CaPJ155), *CSA5*$^{BFA1}$ (CaPJ156) and *CSA6* (CaPJ157), after 8 h of induction with Atc (3 µg/ml) and overnight recovery in a rich medium without Atc. A representative image of an uninduced culture is shown as a reference (leftmost panel, -Atc). Scale bar, 10 µm. **(B)** Representative BFP/GFP density plots of *CSA1*$^{CLB4}$ (CaPJ152), *CSA2*$^{ASE1}$ (CaPJ153) and *CSA3*$^{KIP2}$ (CaPJ154), along with EV (CaPJ150), in presence of Atc (3 µg/ml). **(C)** Mean frequency of CIN ($x10^{-5}$) at the BFP/GFP locus in EV (CaPJ150) versus *CSA1*$^{CLB4}$ (CaPJ152), *CSA2*$^{ASE1}$ (CaPJ153) and *CSA3*$^{KIP2}$ (CaPJ154) overexpression strains; *N*=3. Unpaired *t*-test, one-tailed, *P*-values show a significant difference. **(D)** Analysis of the marker genes, *ARG4*, *HIS1* and *HYGB* by replica plating in $BFP^-$ $GFP^+$ colonies of EV (CaPJ150) and *CSA1*$^{CLB4}$ (CaPJ152), *CSA2*$^{ASE1}$ (CaPJ153) and *CSA3*$^{KIP2}$ (CaPJ154) overexpression strains; *N*=3 with ≥100 colonies for each *N*. **(E)** Cell cycle analysis of EV (CaPJ160) and *CSA4*$^{MCM7}$ (CaPJ165), *CSA5*$^{BFA1}$ (CaPJ166) and *CSA6* (CaPJ167) overexpression strains; *N*=2. Briefly, overnight grown cells were induced for 8 h in presence of Atc (3 µg/ml). Cells were harvested and ethanol-fixed, treated with RNase, stained with propidium iodide and analyzed by flow cytometry for DNA content at specific time intervals. **(F)** *Left*, representative micrographs showing nuclear segregation and mitotic spindle in EV (CaPJ160) and *CSA4*$^{MCM7}$ (CaPJ165), *CSA5*$^{BFA1}$ (CaPJ166) and *CSA6* (CaPJ167) overexpression strains, after 8 h of growth in presence of Dox (50 µg/ml). The nuclear division was analyzed by both Hoechst staining as well as by localization of a KT protein, Cse4-GFP. The spindle integrity was analyzed using Tub4-mCherry, an SPB protein, as a marker. Scale bar, 3 µm. *Right*, quantitation of the cells with indicated phenotypes; *n* ≥100 cells.

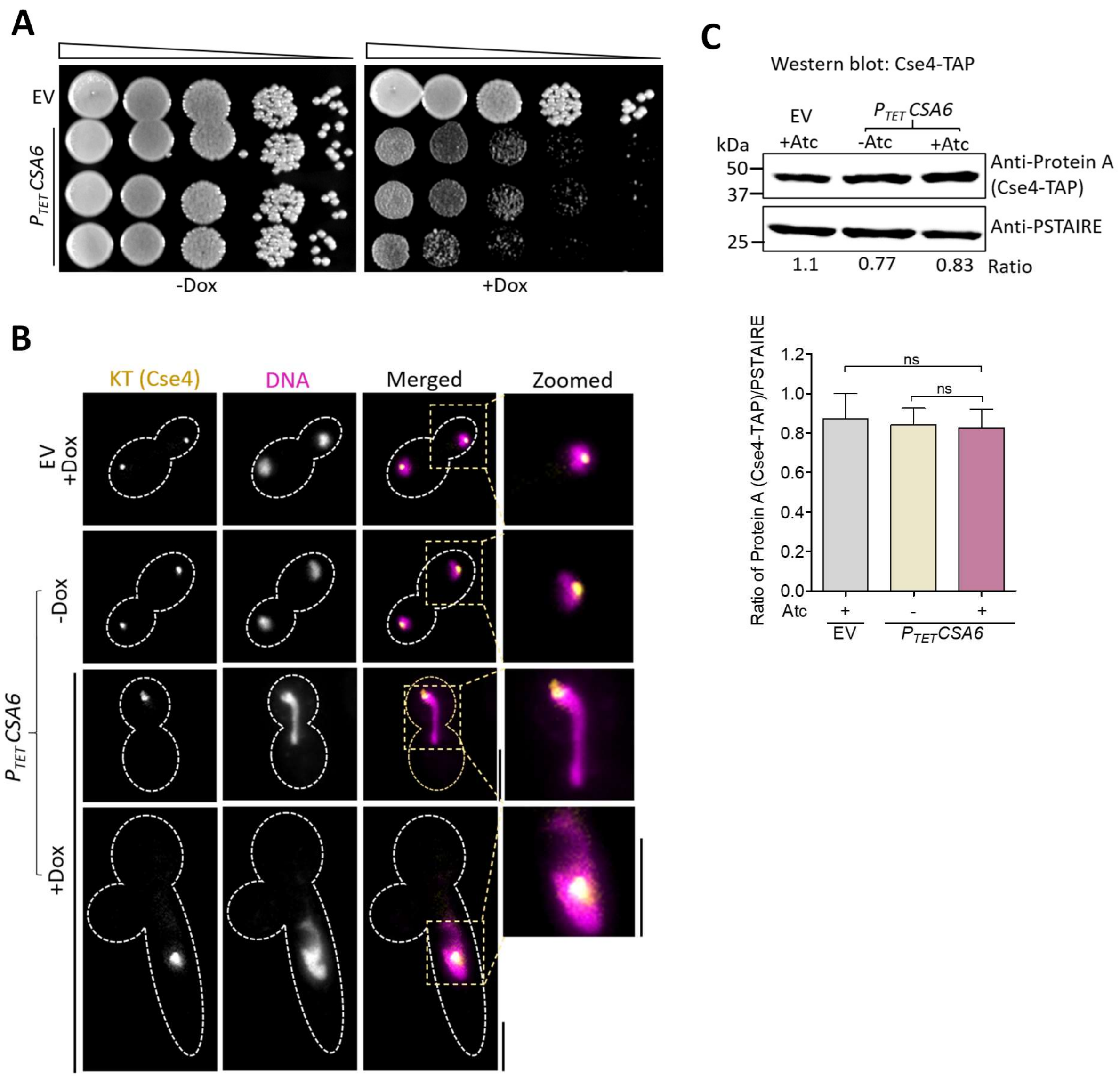


**Fig. S4. Overexpression of *CSA6* affects cell growth but does not perturb kinetochore integrity in *C. albicans*. (A)** Ten-fold serial dilutions, starting from $10^5$ cells, each of CaPJ170 (EV) or CaPJ176 (*CSA6* $^{OE}$) were spotted on YPDU agar plates with or without Dox (50 µg/ml) and incubated at 30°C for two days. **(B)** Localization of Cse4-GFP in CaPJ183 (*CSA6* $^{OE}$ mutant) and CaPJ182 (EV), after 8 h of growth under indicated conditions of Dox (50 µg/ml). The nucleus was stained with Hoechst dye for reference. Scale bar, 5 µm **(C)** *Top*, immunoblot analysis of Cse4-TAP levels in CaPJ173 (EV) and CaPJ179 (*CSA6* $^{OE}$ mutant) in presence or absence of Atc (3 µg/ml) using anti-Protein A antibodies; *N*=3. PSTAIRE was used as a loading control. Cse4-TAP levels were normalized by calculating the ratio of Protein A/PSTAIRE.

*Bottom*, Quantitation of the normalized Cse4-TAP levels; *N*=3. One-way ANOVA and Bonferroni posttest, *P*-values were non-significant (ns) (>0.05).

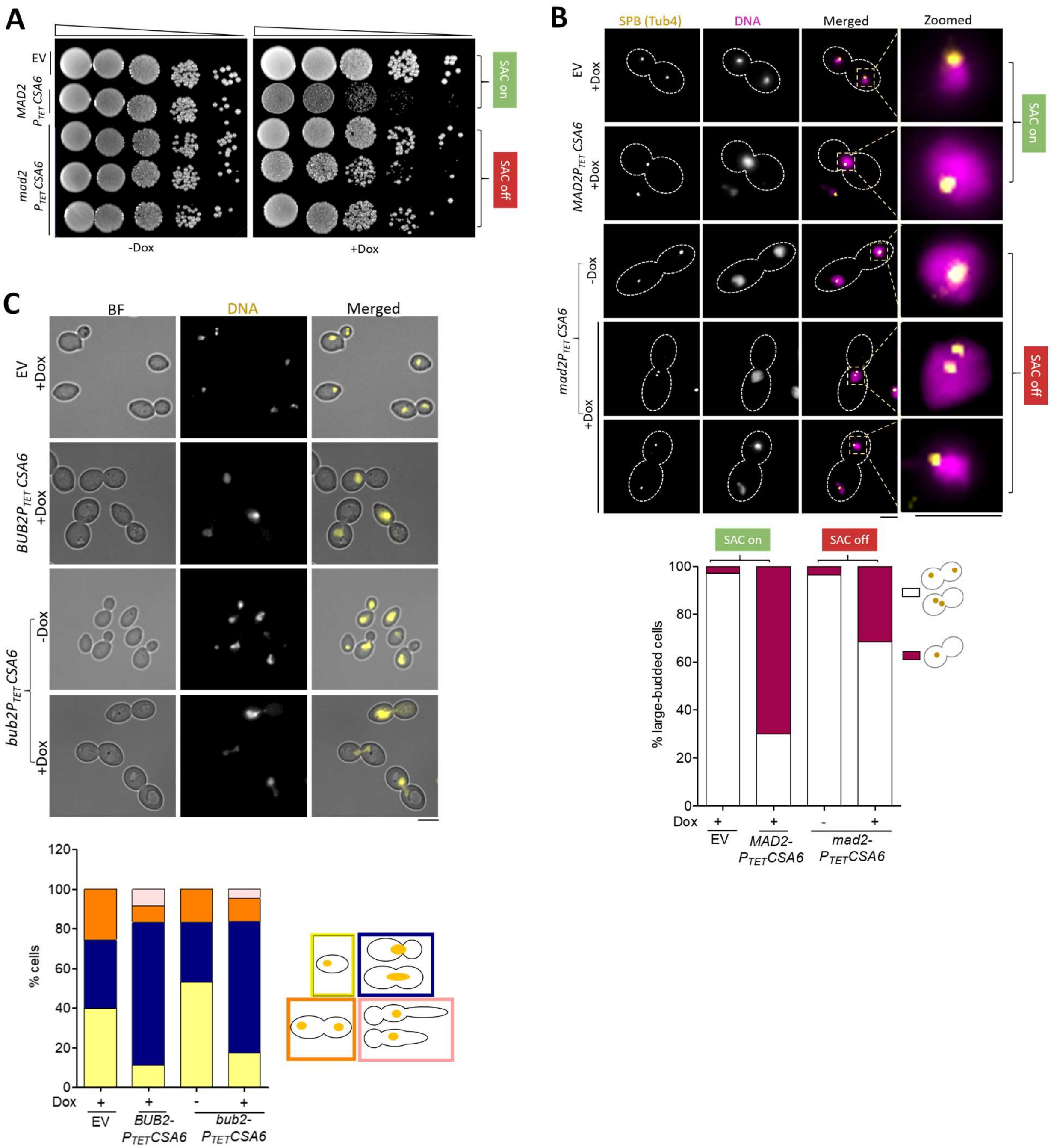


**Fig. S5.** ***CSA6*** **over-expression associated G2/M arrest is relieved upon** ***mad2*** **but not** ***bub2*** **deletion. (A)** Spot dilution analysis of CaPJ170 (EV), CaPJ176 (*MAD2CSA6* $^{OE}$ ) and CaPJ197 (*mad2CSA6*$^{OE}$). Ten-fold serial dilutions, starting from $10^5$ cells, were spotted on YPDU agar plates with or without Dox (50 µg/ml) and incubated at 30°C for two days. **(B)** *Top,* localization patterns of Tub4-GFP in large-budded cells of CaPJ171 (EV), CaPJ177 (*MAD2CSA6*$^{OE}$) and

CaPJ198 (*mad2CSA6* $^{OE}$) after 8 h of growth under indicated conditions of Dox (50 µg/ml). Hoechst staining was done to mark the nuclei. Scale bar, 3 µm. *Bottom*, quantitation of the large-budded cells with the indicated Tub4 phenotypes; *n* ≥100 cells. **(C)** *Top*, representative images of Hoechst-stained CaPJ170 (EV), CaPJ176 (*BUB2CSA6*$^{OE}$) and CaPJ200 (*bub2CSA6*$^{OE}$) after 8 h of growth under indicated conditions of Dox (50 µg/ml). Scale bar, 5 µm. *Bottom*, percent cells with indicated cell types; *n* ≥100 cells.

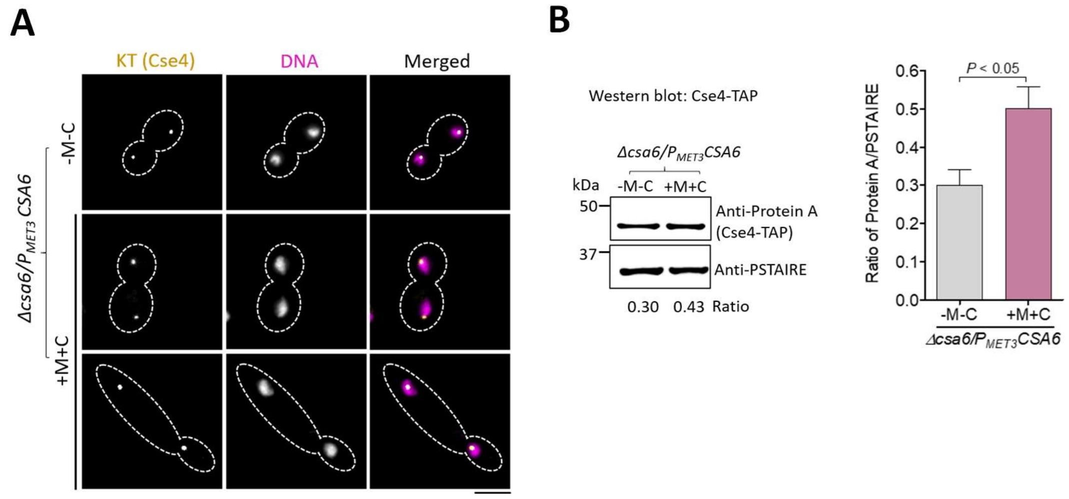


**Fig. S6. Csa6 depleted cells duplicate and segregate their nuclei. (A)** Localization of Cse4-GFP in *CSA6*$^{PSD}$ strain CaPJ213, after 6 h of growth in permissive (YPDU-M-C) or repressive (YPDU + 5 mM M and 5 mM C) conditions. Cse4-GFP colocalized with the nucleus, stained with Hoechst dye. Scale bar, 5 µm. **(B)** *Left*, western blot analysis using anti-Protein A antibodies to compare Cse4-TAP levels in *CSA6*$^{PSD}$ strain CaPJ214 when grown under permissive (YPDU-M-C) or repressive (YPDU + 5 mM M and 5 mM C) conditions for 6 h; *N*=3. PSTAIRE was used as a loading control. Cse4-TAP levels were normalized by calculating the ratio of Protein A/PSTAIRE. *Right*, quantitation of the normalized Cse4 levels; *N*=3. Paired *t*-test, two-tailed, *P*-value shows a significant difference.

**Table S1. Quantification of BFP/GFP loss frequency in EV**

| Sample | Frequency of $BFP^{+}GFP^{-}$ cells ($x10^{-5}$) | Frequency of $BFP^{-}GFP^{+}$ cells ($x10^{-5}$) |
|---|---|---|
| 1 | 7.14 | 19 |
| 2 | 7.52 | 24 |
| 3 | 13 | 16 |
| 4 | 5.7 | 24 |
| 5 | 6.69 | 20 |
| 6 | 7.75 | 15 |
| 7 | 24 | 16 |
| 8 | 18 | 15 |
| 9 | 15 | 22 |
| 10 | 13 | 20 |
| 11 | 15 | 21 |
| 12 | 24 | 19 |
| 13 | 7.8 | 21 |
| 14 | 15 | 22 |
| 15 | 14 | 15 |
| 16 | 11 | 14 |
| 17 | 15 | 13 |
| 18 | 20 | 16 |
| 19 | 14 | 15 |
| 20 | 22 | 20 |
| 21 | 18 | 25 |
| 22 | 15 | 21 |
| Mean | 14.02 | 18.77 |

**Table S2. BFP/GFP loss frequency in the primary hits**

| ORF no. | Orthologs in *S. cerevisiae* | Frequency of $BFP^{+}GFP^{-}$ cells ($x10^{-5}$) | Fold change | Frequency of $BFP^{-}GFP^{+}$ cells ($x10^{-5}$) | Fold change |
|---|---|---|---|---|---|
| 19.1447 | - | 180 | 13.0 | 210 | 11.2 |
| 19.7186 | *CLB4* | 180 | 13.0 | 180 | 9.6 |
| 19.608 | *BFA1* | 140 | 10.0 | 190 | 10.2 |
| 19.3135 | *UBX2* | 120 | 8.6 | 45 | 2.4 |
| 19.202 | *MCM7* | 82 | 5.9 | 120 | 6.4 |
| 19.1048 | *IFD6* | 63 | 4.5 | 83 | 4.4 |
| 19.6588 | *NBP2* | 83 | 5.9 | 40 | 2.1 |
| 19.1601 | *RPL3* | 70 | 5.0 | 43 | 2.3 |
| 19.3437 | - | 72 | 5.1 | 37 | 2.0 |
| 19.1934 | *HST3* | 66 | 4.7 | 39 | 2.0 |
| 19.1542 | *HEX3* | 50 | 3.6 | 54 | 2.9 |
| 19.6778 | *DRS2* | 43 | 3.0 | 55 | 2.9 |
| 19.4153 | *ULA1* | 52 | 3.7 | 42 | 2.2 |
| 19.1396 | *AGE2* | 53 | 3.8 | 39 | 2.0 |
| 19.3349 | *RPB2* | 38 | 2.7 | 44 | 2.3 |
| 19.1747 | *KIP2* | 41 | 2.9 | 50 | 2.7 |
| 19.4979 | *KNS1* | 40 | 2.8 | 55 | 2.9 |
| 19.7377 | *ASE1* | 40 | 2.9 | 48 | 2.6 |
| 19.1999 | - | 33 | 2.3 | 44 | 2.3 |
| 19.3421.1 | *ROX3* | 35 | 2.5 | 47 | 2.5 |
| 19.6118 | *DSS4* | 38 | 2.7 | 39 | 2.1 |
| 19.4340.1 | *SMX3* | 33 | 2.4 | 39 | 2.1 |
| 19.5212 | *CST9* | 32 | 2.3 | 36 | 1.9 |

**Table S3. Strains used in this study**

| Name (Description) | Genotype | Reference |
|---|---|---|
| SN148 | *Δura3::imm434/Δura3::imm434, Δhis1::hisG/Δhis1::hisG, Δarg4::hisG/Δarg4::hisG, Δleu2::hisG/Δleu2::hisG* | (*19*) |
| YJB8675 | *Δura3::imm434/Δura3::imm434, Δhis1::hisG/Δhis1::hisG, Δarg4::hisG/Δarg4::hisG, CSE4-GFPCSE4/CSE4* | (*20*) |
| J110 | SN148 *mad2::ARG4/mad2::LEU2* | (*14*) |
| CEC3867 | SN148 *Ca21ch4_C_albicans_SC5314:473390 to 476401Δ::PTDH3-GFP-ARG4/Ca21ch4_C_albicans_SC5314:473390 to 476401Δ::PTDH3-BFP-HIS1, ADH1/adh1::PTDH3-cartTA-SAT1* | (*5*) |
| CAKS102 | SN148 *CSE4/CSE4-TAP::URA3* | (*13*) |
| Cd36 (*C. dubliniensis* prototroph) | *URA3/URA3* (clinical isolate) | (*12*) |
| CEC5201 (CSA reporter) | CEC3867 *Ca22ch4_C_albicans_SC5314:1452840 to 1453029*Δ*::*$P_{TDH3}$*-RFP-Hyg*$^{R}$*/Ca22ch4_C_albicans_SC5314:1452840 to 1453029* | This study |
| CaPJ148 (Mono-BFP) | *SN148 Ca21ch4_C_albicans_SC5314:473390 to 476401Δ::PTDH3-BFP-HIS1/Ca21ch4_C_albicans_SC5314:473390 to 476401* | This study |
| CaPJ149 (Mono-GFP) | *SN148 Ca21ch4_C_albicans_SC5314:473390 to 476401*Δ*::PTDH3-GFP-ARG4/Ca21ch4_C_albicans_SC5314:473390 to 476401* | This study |
| CaPJ150 (EV in CSA reporter) | CEC5201 *RPS1/RPS1::PTET-GtwB-URA3* | This study |
| CaPJ151 (*CDC20*$^{OE}$ in CSA reporter) | CEC5201 *RPS1/RPS1::PTET-CDC20-URA3* | This study |
| CaPJ152 (*CSA1*$^{CLB4}$ overexpression in CSA reporter) | CEC5201 *RPS1/RPS1::PTET-CLB4-URA3* | This study |
| CaPJ153 (*CSA2*$^{ASE1}$ overexpression in CSA reporter) | CEC5201 *RPS1/RPS1::PTET-ASE1-URA3* | This study |
| CaPJ154 (*CSA3*$^{KIP2}$ | CEC5201 *RPS1/RPS1::PTET-KIP2-URA3* | This study |

| | | |
|---|---|---|
| overexpression in CSA reporter) | | |
| CaPJ155 (*CSA4*$^{MCM7}$ overexpression in CSA reporter) | CEC5201 *RPS1/RPS1::PTET-MCM7-URA3* | This study |
| CaPJ156 (*CSA5*$^{BFA1}$ overexpression in CSA reporter) | CEC5201 *RPS1/RPS1::PTET-BFA1-URA3* | This study |
| CaPJ157 (*CSA6*$^{OE}$ in CSA reporter) | CEC5201 *RPS1/RPS1::PTET-CSA6-URA3* | This study |
| CaPJ158 | YJB8675 *ADH1/adh1::PTDH3-cartTA-SAT1* | This study |
| CaPJ159 | CaPJ158 *TUB4/TUB4-mCherry::ARG4* | This study |
| CaPJ160 (EV in *CSE4-GFP, TUB4-mCherry*) | CaPJ159 *RPS1/RPS1::PTET-GtwB-URA3* | This study |
| CaPJ165 (*CSA4*$^{MCM7}$ overexpression in *CSE4-GFP, TUB4-mCherry*) | CaPJ159 *RPS1/RPS1::PTET-MCM7-URA3* | This study |
| CaPJ166 (*CSA5*$^{BFA1}$ overexpression in *CSE4-GFP, TUB-4mCherry*) | CaPJ159 *RPS1/RPS1::PTET-BFA1-URA3* | This study |
| CaPJ167 (*CSA6*$^{OE}$ in *CSE4-GFP, TUB4-mCherry*) | CaPJ159 *RPS1/RPS1::PTET-CSA6 -URA3* | This study |
| CaPJ169 | SN148 *ADH1/adh1::PTDH3-cartTA-SAT1* | This study |
| CaPJ180 (*CSA6-TAP*) | SN148 *CSA6/CSA6-TAP::ARG4* | This study |
| CaPJ181 (*PTETCSA6-TAP*) | CaPJ169 *RPS1/RPS1::PTET-CSA6-TAP-URA3* | This study |
| CaPJ170 (EV in SN148) | CaPJ169 *RPS1/RPS1::PTET-GtwB-URA3* | This study |
| CaPJ176 (*CSA6*$^{OE}$ in SN148) | CaPJ169 *RPS1/RPS1::PTET-CSA6-URA3* | This study |
| CaPJ162 | YJB8675 *TUB1/TUB1-mCherry::HIS1* | This study |
| CaPJ163 | CaPJ162 *ADH1/adh1::PTDH3-cartTA-SAT1* | This study |

| | | |
|---|---|---|
| CaPJ182 (EV in *CSE4-GFP*) | CaPJ163 *RPS1/RPS1::PTET-GtwB-URA3* | This study |
| CaPJ183 (*CSA6*$^{OE}$ in *CSE4-GFP*) | CaPJ163 *RPS1/RPS1::PTET-CSA6-URA3* | This study |
| CaPJ173 (*CSE4-TAP* in EV) | CaPJ170 *CSE4/CSE4-TAP::LEU2* | This study |
| CaPJ179 (*CSE4-TAP* in *CSA6*$^{OE}$) | CaPJ176 *CSE4/CSE4-TAP::LEU2* | This study |
| CaPJ171 (EV in *TUB4-GFP*) | CaPJ170 *TUB4/TUB4-GFP:: HIS1* | This study |
| CaPJ172 (EV in *TUB4-GFP, TUB1-mCherry*) | CaPJ170 *TUB4/TUB4-GFP:: HIS1, TUB1/TUB1-mCherry::ARG4* | This study |
| CaPJ177 (*CSA6*$^{OE}$ in *TUB4-GFP*) | CaPJ176 *TUB4/TUB4-GFP:: HIS1* | This study |
| CaPJ178 (*CSA6*$^{OE}$ in *TUB4-GFP, TUB1-mCherry*) | CaPJ177 *TUB1/TUB1-mCherry::ARG4* | This study |
| CaPJ196 | CaJ110 *ADH1/adh1::PTDH3-cartTA-SAT1* | This study |
| CaPJ197 (*CSA6*$^{OE}$ in *mad2*) | CaPJ196 *RPS1/RPS1::PTET-CSA6-URA3* | This study |
| CaPJ198 (*CSA6*$^{OE}$ in *mad2, TUB4-GFP)* | CaPJ197 *TUB4/TUB4-GFP:: HIS1* | This study |
| CaPJ109 | SN148 *bub2::FRT/BUB2* | This study |
| CaPJ110 | SN148 *bub2::FRT/ bub2::FRT* | This study |
| CaPJ199 | CaPJ110 *ADH1/adh1::PTDH3-cartTA-SAT1* | This study |
| CaPJ200 (*CSA6*$^{OE}$ in *bub2*) | CaPJ199 *RPS1/RPS1::PTET-CSA6-URA3* | This study |
| CaPJ209 (*CSA6* heterozygous null in SN148) | SN148 *csa6::FRT/CSA6* | This study |
| CaPJ210 (*CSA6*$^{PSD}$ in SN148) | SN148 *csa6::FRT/MET3prCSA6::URA3* | This study |
| CaPJ212 ($P_{MET3}$*CSA6-TAP*) | SN148 *csa6::FRT/MET3prCSA6-TAP-ARG4::URA3* | This study |
| CaPJ113 | YJB8675 *csa6::FRT/CSA6* | This study |

| | | |
|---|---|---|
| CaPJ213 (*CSA6^PSD* in *CSE4-GFP*) | YJB8675 *csa6::FRT/MET3prCSA6::URA3* | This study |
| CaPJ214 (*CSE4-TAP* in *CSA6^PSD*) | CaPJ210 *CSE4/CSE4-TAP::LEU2* | This study |
| CaPJ211 (*CSA6^PSD* in *TUB4-GFP, TUB1-mCherry*) | CaPJ210, *TUB4/TUB4-GFP:: HIS1, TUB1/TUB1-mCherry::ARG4* | This study |
| CaPJ216 (*SOL1-TAP*) | SN148 *SOL1/SOL1-TAP::HIS1* | This study |
| CaPJ217 ($P_{TET}$*SOL1-TAP*) | CaPJ169 *RPS1/RPS1::PTET-SOL1-TAP-URA3* | This study |
| CaPJ215 (*CSA6^PSD* in *SOL1^OE*) | SN148 *csa6::FRT/MET3prCSA6::HIS1, ADH1/adh1::PTDH3-cartTA-SAT1, RPS1/RPS1::PTET-SOL1-URA3* | This study |
| CaPJ218 (*TEM1-GFP* in *CSA6^PSD*) | SN148 *csa6::FRT/MET3prCSA6::URA3, TUB4/TUB4-mCherry::NAT, TEM1/TEM1-GFP::HIS1* | This study |
| CaPJ119 (*CSA6-mCherry* in *CSE4-GFP*) | YJB8675 *CSA6/CSA6-mCherry::ARG4* | This study |
| CaPJ117 | SN148 *csa6::FRT/ CSA6-mCherry::ARG4* | This study |
| CaPJ118 | SN148 *CSA6/CSA6-mCherry::ARG4* | This study |
| CaPJ120 (*CSA6-mCherry* in *TUB4-GFP*) | CaPJ118 *TUB4/TUB4-GFP::HIS1* | This study |
| CaPJ121 (*CSA6-mCherry* in *SPC110-GFP*) | CaPJ118 *SPC110/SPC110-GFP::HIS1* | This study |
| CaPJ300 | CaPJ209 *TUB4/TUB4-mCherry::ARG4 + pCdCSA6-GFP-ARS2::URA3* | This study |
| CaPJ301 | SN148 *csa6::FRT/MET3prCSA6::HIS1, TUB4/TUB4-mCherry::ARG4* | This study |
| CaPJ302 | SN148 *csa6::FRT/MET3prCSA6::HIS1, TUB4/TUB4-mCherry::ARG4 + pCdCSA6-GFP-ARS2::URA3* | This study |

**Table S4. Primers used in this study**

| Name | Sequence | Description |
|---|---|---|
| RFP-PstI-F | AAACCCctgcagAAAGATGGTTTCTAAAGGTG | RFP-HygB K7 |
| RFP-NheI-R | CCCAAAgctagcCATATTATTATCTTCAGAAG | RFP-HygB K7 |
| K7_BFP_GFP_Ch4_Right_F | TATATATTTCTGGCAATGCAGCAATTCTCGGATATCACCGAAAAAAAAGATCTTAGCGGGCACGACACGACTCTCTTGATATAAGCGAATTTTCAGTATCAGGAAACAGCTATGACC | Integration of the RFP-HygB K7 on the right arm of Chr4 |
| RFP_Insertion_Ch4_Right_Reverse | TCTCTATACGAGTTAAGAGTAGTCTTACAATAGTCTATAGATAGAATTTCAGACCTTTTTGTGTGGGTATTGCCGAAATTCTTTTTCCAGAAGATGACGAGAGAAAATACCCGTGACG | Integration of the RFP-HygB K7 on the right arm of Chr4 |
| PJ86 | ACAAGCTTATTGAGTGACGAAAAGTC | Confirmation of pNIMX integration |
| PJ87 | TTTACGGGTTGTTAAACCTTCGATTC | |
| PJ88 | ATACTACTGAAAATTTCCTGACTTTC | Confirmation of overexpression plasmid integration |
| PJ89 | ATTACTATTTACAATCAAAGGTGGTC | |
| PJ90 | ATCAACAAGTTTGTACAAA | Sequencing of overexpression plasmid |
| SR149 | CgcACTAGTATGGTTTCAAAAGGTGAAGAAG | Amplification of mCherry-coding gene |
| SR150 | ggaCCCGGGACCCAGAAAGCATTCATCGCG | |
| LS39FP | TCCCCGCGGGATCGATATAAAACTAATCGTGTTAG | C-term tagging of Tub4 with GFP/mCherry |
| LS39RP | GGACTAGTTATACCCATATCTGCATCATCTATATTG | |
| PJ77 | tataCCGCGGACTGTTCAATTAGTCGATTGGTGTC | C-term tagging of Tub1 with mCherry |
| PJ78 | atatACTAGTATATTCTTCTTCTTCTTCAGGGAAAG | |
| NV241 | TAA GGG CCC CCA GCT GCT ACT TCC TC | C-term tagging of Cse4 with TAP |
| NV242 | acgc GTCGAC GGCCAATTATAAATGTGAAGGG | |
| PJ108 | atatAGATCTAATAAGAATACGCTATCTCC | C-term tagging of Csa6 with TAP |
| PJ109 | atatTTAATTAAGTTAGAACGACCCATAAATTC | |
| NV34 | GAGCACGTATTGGGTTTGC | Confirmation of TAP cassette integration |
| PJ127 | atatGATATCATGGAAGATTCAACTGAAGATATAATTA | Cloning of *CSA6TAP* under $P_{TET}$ |
| PJ128 | atatGATATCTCACTGATGATTCGCGTC | |

| | | |
|---|---|---|
| PJ110 | atatGGTACCAATGCTAGTAGGGTCTAGAC | Deletion cassette for *BUB2* |
| PJ111 | atatCTCGAGTTTGTATCGGAAGGATGTAG | |
| PJ112 | atatCCGCGGATCAATTCTGCACATGGTATG | |
| PJ113 | atatGAGCTCTTGCCTAATAAGACGCCAATTC | |
| PJ114 | atatCCGCGGATATACGCTTTCCCTTCG | |
| PJ115 | atatGAGCTCAATTCTTTAGGAACTTTTCTATCG | |
| PJ116 | AGTCTTGAACGAAAAAGTCTAG | |
| PJ3 | CTATTCTCTAGAAAGTATAGGAACTTC | Confirmation of pPSFS2a integration |
| PJ118 | acgcctaacatatgtgaagtg | |
| PJ95 | atatGGTACCAAGAAGCATGTGGTATGAAGCAC | Deletion cassette for *CSA6* |
| PJ96 | atatCTCGAGTTGTGTCGGTCTGTACGTG | |
| PJ97 | atatCCGCGGGTGGGTAGGTTACACAGAGTC | |
| PJ98 | atatGAGCTCTGGTCCACTACAACCCCTTTTG | |
| PJ99 | TGGCTGATATGGCTCATTG | |
| PJ93 | atgtGGATCCATGGAAGATTCAACTGAAGATATAATTA | Cloning of *CSA6* under *MET3* |
| PJ94 | atctCTGCAGATTATATGCAGCAGATTGAGAAGG | |
| PJ141 | atatGGATCCATAACTCTTTCACGCAAGCTC | C-term tagging of Sol1 with TAP |
| PJ142 | atatTTAATTAATATATTATCAAACGATAATCTCTTTGGTTTG | |
| PJ119 | atatGATATCATGTCCTCTTCTAATGATACACCATC | Cloning of *SOL1* under $P_{TET}$ |
| PJ120 | atatGATATCTTATATATTATCAAACGATAATCTCTTTGGTTTG | |
| PJ121 | atatCCGCGGACAAGAAAGTCTACGCTAAATTC | C-term tagging of Tem1 with GFP |
| PJ122 | atatACTAGTCTTATATATCAATATGGGTTCCCCCAC | |
| PJ123 | TGCTACCATTGGTCTCAAATGATG | |
| PJ124 | ccatacgcgaaagtagtg | Confirmation of GFP cassette integration |
| TEJ1 | attaCCGCGGAATAAGAATACGCTATCTCC | C-term tagging of Csa6 with mCherry |
| TEJ2 | atgcACTAGTGTTAGAACGACCCATAAATTC | |
| TEJ13 | TCGAAGAAATGCTGTCC | |
| TEJ14 | TCTTCTTCACCTTTTGAAACC | Confirmation of mCherry cassette integration |

| | | |
|---|---|---|
| PJ106 | atatCCGCGGAATTGAAGAAAGAGGTTCAAGAC | C-term tagging of Spc110 with GFP |
| PJ107 | atatACTAGTATTGTATTTAAGTCTGGCCAC | |
| VS5 | AGTCTCTAGACAAGTATTCAACAATTTCTGTC | Ectopic expression of *C. dubliniensis* Csa6, tagged with GFP |
| VS6 | GTGAAAAGTTCTTCTCCCTTACTCATATTGGAATGGCCCATAAATTCTG | |
| VS7 | CAGAATTTATGGGCCATTCCAATATGAGTAAGGGAGSSGAACTTTTCAC | |
| VS8 | AGTCCTGCAGGGGCATTTTATGATGGAATGAATG | |

**Table S5. Plasmids used in this study**

| Name | Description | Reference |
|---|---|---|
| CIp10-P$_{TET}$-GTW derivatives | Overexpression plasmid collection | (*6*) |
| pNIMX | Plasmid harbouring P$_{TET}$ transactivator | (*1*) |
| pGFP-HIS | GFP-tagging plasmid | (*9*) |
| pRFP-Arg4 | mCherry-tagging plasmid | (*10*) |
| pFA-TAP-*ARG4* | TAP-tagging plasmid | (*17*) |
| pFA-TAP-*HIS1* | | |
| pSFS2a | Recyclable *SAT1*-flipper cassette | (*15*) |
| pCaDis | Plasmid for promoter replacement with *MET3*pr | (*16*) |
| pBSNAT | Plasmid used for cloning *NAT1* | (*12*) |
| pCaADH1-yEmRFP | Plasmid used for cloning mCherry | (*11*) |
| pMad2-2 | Plasmid used for cloning *CSE4-TAP* fragment | (*14*) |
| pNIM1R-RFP | Plasmid used for cloning RFP | (*2*) |
| pTDH3-GFP-URA3 | Plasmid used for generating the pTDH3-RFP-HygB plasmid | (*3*) |
| pAU34-CaHygB | Plasmid used for cloning HygB | (*4*) |
| pTub4-GFP-His | GFP-tagging plasmid for Tub4 | This study |
| pTub4-mCherry-Arg4 | mCherry-tagging plasmids for Tub4 | This study |
| pTub4-mCherry-Nat | | This study |
| pTub1-mCherry-Arg4 | mCherry-tagging plasmid for Tub1 | This study |
| pCse4-TAP-Leu | TAP-tagging plasmid for Cse4 | This study |
| pCsa6-TAP-Arg | TAP-tagging plasmid for Csa6 | This study |
| CIp10-P$_{TET}$-Csa6TAP | Overexpression plasmid for *CSA6-TAP* | This study |
| pBub2del#1 | Deletion cassettes for *BUB2* | This study |
| pBub2del#2 | | This study |
| pCsa6del | Deletion cassette for *CSA6* | This study |
| pCsa6-Met3-Ura | Plasmids for promoter replacement of *CSA6* with *MET3* | This study |
| pCsa6-Met3-His | | This study |
| CIp10-P$_{TET}$-SOL1 | Overexpression plasmid for *SOL1* | This study |
| CIp10-P$_{TET}$-SOL1TAP | Overexpression plasmid for *SOL1-TAP* | This study |
| pTEM1-GFP-His | GFP-tagging plasmid for Tem1 | This study |
| pCsa6-mCherry-Arg | mCherry-tagging plasmid for Csa6 | This study |
| pSpc110-GFP-His | GFP-tagging plasmid for Spc110 | This study |
| pCdCsa6-GFP-ARS2 | Ectopic expression of GFP-tagged *C. dubliniensis* Csa6 | This study |

### References


1. M. Chauvel, A. Nesseir, V. Cabral, S. Znaidi, S. Goyard, S. Bachellier-Bassi, A. Firon, M. Legrand, D. Diogo, C. Naulleau, T. Rossignol, C. d'Enfert, A versatile overexpression strategy in the pathogenic yeast *Candida albicans*: identification of regulators of morphogenesis and fitness. *PLoS One* **7**, e45912 (2012).
2. D. Prieto, E. Roman, I. Correia, J. Pla, The HOG pathway is critical for the colonization of the mouse gastrointestinal tract by *Candida albicans*. *PLoS One* **9**, e87128 (2014).
3. S. Znaidi, L. van Wijlick, A. Hernandez-Cervantes, N. Sertour, J. L. Desseyn, F. Vincent, R. Atanassova, V. Gouyer, C. A. Munro, S. Bachellier-Bassi, F. Dalle, T. Jouault, M. E. Bougnoux, C. d'Enfert, Systematic gene overexpression in *Candida albicans* identifies a regulator of early adaptation to the mammalian gut. *Cell Microbiol.* **20**, e12890 (2018).
4. L. R. Basso, Jr., A. Bartiss, Y. Mao, C. E. Gast, P. S. Coelho, M. Snyder, B. Wong, Transformation of *Candida albicans* with a synthetic hygromycin B resistance gene. *Yeast* **27**, 1039-1048 (2010).
5. A. Feri, R. Loll-Krippleber, P. H. Commere, C. Maufrais, N. Sertour, K. Schwartz, G. Sherlock, M. E. Bougnoux, C. d'Enfert, M. Legrand, Analysis of Repair Mechanisms following an Induced Double-Strand Break Uncovers Recessive Deleterious Alleles in the *Candida albicans* Diploid Genome. *mBio* **7**, (2016).
6. M. Legrand, S. Bachellier-Bassi, K. K. Lee, Y. Chaudhari, H. Tournu, L. Arbogast, H. Boyer, M. Chauvel, V. Cabral, C. Maufrais, A. Nesseir, I. Maslanka, E. Permal, T. Rossignol, L. A. Walker, U. Zeidler, S. Znaidi, F. Schoeters, C. Majgier, R. A. Julien, L. Ma, M. Tichit, C. Bouchier, P. Van Dijck, C. A. Munro, C. d'Enfert, Erratum: Generating genomic platforms to study *Candida albicans* pathogenesis. *Nucleic Acids Res.* **46**, 8664 (2018).
7. A. J. Harwood, The rapid boiling method for small-scale preparation of plasmid DNA. *Methods Mol. Biol.* **58**, 265-267 (1996).
8. A. Walther, J. Wendland, An improved transformation protocol for the human fungal pathogen *Candida albicans*. *Curr. Genet.* **42**, 339-343 (2003).

9. G. Chatterjee, S. R. Sankaranarayanan, K. Guin, Y. Thattikota, S. Padmanabhan, R. Siddharthan, K. Sanyal, Repeat-Associated Fission Yeast-Like Regional Centromeres in the Ascomycetous Budding Yeast *Candida tropicalis*. *PLoS Genet.* **12**, e1005839 (2016).
10. N. Varshney, K. Sanyal, Aurora kinase Ipl1 facilitates bilobed distribution of clustered kinetochores to ensure error-free chromosome segregation in *Candida albicans*. *Mol. Microbiol.* **112**, 569-587 (2019).
11. S. Keppler-Ross, C. Noffz, N. Dean, A new purple fluorescent color marker for genetic studies in *Saccharomyces cerevisiae* and *Candida albicans*. *Genetics* **179**, 705-710 (2008).
12. J. Thakur, K. Sanyal, Efficient neocentromere formation is suppressed by gene conversion to maintain centromere function at native physical chromosomal loci in *Candida albicans*. *Genome Res.* **23**, 638-652 (2013).
13. S. Mitra, J. Gomez-Raja, G. Larriba, D. D. Dubey, K. Sanyal, Rad51-Rad52 mediated maintenance of centromeric chromatin in *Candida albicans*. *PLoS Genet.* **10**, e1004344 (2014).
14. J. Thakur, K. Sanyal, The essentiality of the fungus-specific Dam1 complex is correlated with a one-kinetochore-one-microtubule interaction present throughout the cell cycle, independent of the nature of a centromere. *Eukaryot. Cell* **10**, 1295-1305 (2011).
15. O. Reuss, A. Vik, R. Kolter, J. Morschhauser, The SAT1 flipper, an optimized tool for gene disruption in *Candida albicans*. *Gene* **341**, 119-127 (2004).
16. R. S. Care, J. Trevethick, K. M. Binley, P. E. Sudbery, The MET3 promoter: a new tool for *Candida albicans* molecular genetics. *Mol. Microbiol.* **34**, 792-798 (1999).
17. H. Lavoie, A. Sellam, C. Askew, A. Nantel, M. Whiteway, A toolbox for epitope-tagging and genome-wide location analysis in *Candida albicans*. *BMC Genomics* **9**, 578 (2008).
18. R. D. Cannon, H. F. Jenkinson, M. G. Shepherd, Isolation and nucleotide sequence of an autonomously replicating sequence (ARS) element functional in *Candida albicans* and *Saccharomyces cerevisiae*. *Mol. Gen. Genet.* **221**, 210-218 (1990).
19. S. M. Noble, A. D. Johnson, Strains and strategies for large-scale gene deletion studies of the diploid human fungal pathogen *Candida albicans*. *Eukaryot. Cell* **4**, 298-309 (2005).

20. A. P. Joglekar, D. Bouck, K. Finley, X. Liu, Y. Wan, J. Berman, X. He, E. D. Salmon, K. S. Bloom, Molecular architecture of the kinetochore-microtubule attachment site is conserved between point and regional centromeres. *J. Cell Biol.* **181**, 587-594 (2008).
